# High-throughput selection of glucose-binding proteins from massive datasets: Integrating molecular docking and molecular dynamics simulations

**DOI:** 10.1101/2024.03.20.585966

**Authors:** Anurag Makare, Amit Chaudhary, Debankita De, Parijat Deshpande, Ajay Singh Panwar

## Abstract

Selecting suitable glucose-binding proteins (GBPs) is vital for biosensor development for medical diagnostics and quality control in the food industry. Biosensors offer advantages such as high specificity, selectivity, fast response time, continuous measurement, and cost-effectiveness. The current work utilized a combination of molecular docking, molecular dynamics (MD) simulations, and free energy calculations to develop a high-throughput bioinformatics pipeline to select GBP candidates from an extensive protein database (37,325 proteins). Using molecular docking, GBPs with good binding affinity to glucose (1,447 candidates) were virtually screened from the Protein Data Bank. MD simulations ascertained the binding dynamics of a few selected candidates. Further, steered MD (Brownian dynamics fluctuation-dissipation-theorem) was used to estimate binding free energies of the ligand-protein complex. Correlations between ligand-binding parameters obtained from longer MD simulations and binding parameters interpreted from significantly faster docking simulations were investigated. The correlation plots suggested that a combination of threshold values of the following three docking parameters: docking binding energy, binding cavity depth, and the number of hydrogen bonds between the ligand and binding site residues can be used to predict candidate GBPs reliably. Thus, a high-throughput and accurate protein selection process based on relatively faster docking simulations was proposed to screen GBPs for glucose biosensing.

## 1. Introduction

Biosensors have revolutionized a wide range of fields, from medical diagnostics to environmental monitoring, by combining the specificity of biomolecular recognition with the sensitivity of modern sensing technologies[1]. These powerful devices enable real-time and accurate detection of analytes in complex biological or environmental samples. Proteins are versatile candidates for recognition elements in biosensors, offering unique binding capabilities and adaptability[2–5]. Glucose biosensing is crucial in medical diagnostics and blood glucose monitoring. In numerous applications, such as diabetes management, food industry quality control, and fermentation processes, the precise measurement of glucose is of paramount importance due to its role as an essential energy source[4,6]. Glucose biosensors employ specific enzymes for accurate glucose measurement. Continuous glucose monitoring systems provide real-time data wirelessly for monitoring or smartphones [7–9]. Non-invasive glucose biosensors offer less invasive monitoring through optical and electrochemical methods. These advancements enhance diabetes management, promising ongoing improvements in accuracy and convenience[10–15].

To achieve high selectivity and sensitivity in glucose sensing, selecting an appropriate recognition element becomes critical. In this context, GBPs are promising candidates owing to their innate ability to bind glucose molecules selectively. GBPs encompass a diverse group of proteins with distinct binding pockets or domains that exhibit an affinity for glucose. As recognition elements in biosensors, GBPs offer advantages of high binding affinity, specificity, and stability. Moreover, their potential for genetic and protein engineering allows for performance enhancement and customization to specific applications. The screening and selection of suitable GBPs for biosensor development constitute crucial steps in harnessing their potential. To this end, advanced screening techniques, including combinatorial libraries, phage display, and high-throughput screening methods[16–18], have facilitated the identification of novel GBPs with enhanced binding properties. These techniques enable the exploration of vast protein libraries and the identification of GBPs with desired attributes, such as high affinity, selectivity, and resistance to interfering substances.

Molecular docking enables the analysis of protein-substrate interactions, the assessment of substrate specificity, the evaluation of docking scores, and the optimization of enzyme structures. By predicting binding affinities and stability, molecular docking aids in identifying enzymes with strong substrate binding and high selectivity [19–22]. This computational approach significantly enhances the efficiency of enzyme selection, enabling the development of sensitive and specific biosensors for diverse applications. Additionally, MD simulation of docked complexes provides insights into the dynamic behavior of enzyme-substrate interactions, further elucidating the stability and conformational changes crucial for biosensor design and optimization. In the current work, a bioinformatics pipeline was used to screen glucose-specific GBPs from large datasets of proteins acquired from a protein database. Sequence-based analysis was done to filter out unique GBPs, and molecular docking and MD simulation were employed to screen specific GBPs. A unique set of GBPs with high binding affinity to the glucose molecule were identified.

## 2. Methodology

The overall methodology employed in this study is illustrated in Figure 1. A combination of computational tools was utilized to screen for GBPs.

**Figure 1:**
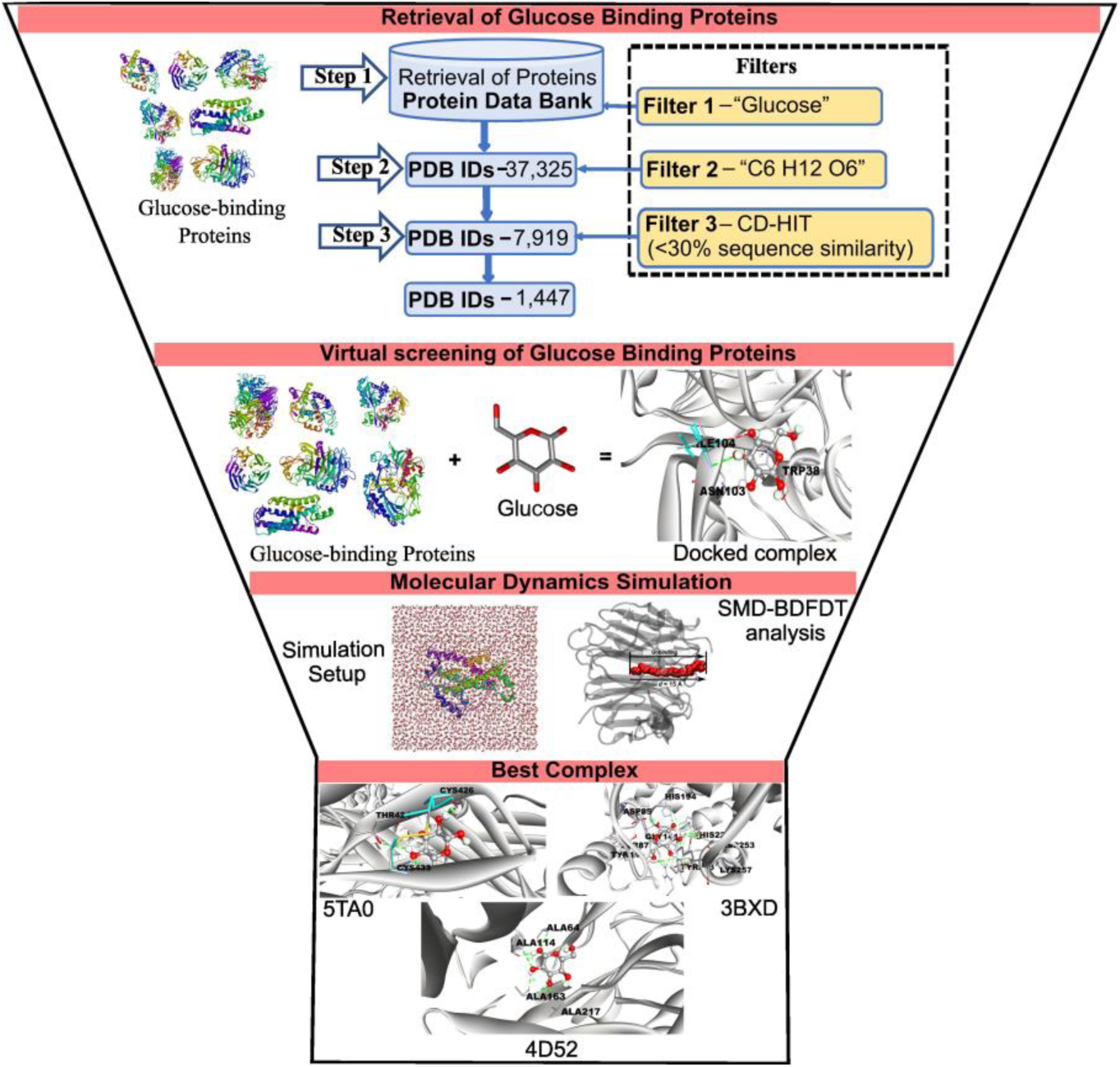
Schematic of the bioinformatics pipeline used to screen the best candidate glucose-binding proteins (GBPs). The proposed scheme is based on a combination of molecular docking and MD simulations.

### 2.1 Virtual screening of glucose-binding proteins (GBPs)

The 3D structure of glucose (PubChem ID 5793) was retrieved from the PubChem database and subsequently prepared by AutoDock 4 for docking against filtered GBPs. In the study, glucose-binding proteins were downloaded from the PDB database [23] using the keyword “glucose” and then filtered out by using two filters (Glucose & Homology filter). Again, AutoDock 4 was employed for protein preparation for molecular docking. AutoDock Vina [24] was employed for protein-ligand docking. The Parallelized Open Babel & AutoDock Suite Pipeline (POAP) [25] facilitated the automation of docking for multiple proteins and ligands. Discovery Studio Visualizer was used to visualize protein-ligand interactions [26].

### 2.2 Filtering and elimination of redundant GBPs

Several filters were implemented to optimize computational time and cost to eliminate redundant proteins. Redundancy criteria encompassed proteins unrelated to glucose, mutations within the same proteins, and homologous proteins. The integration of these filters facilitated the refinement of our dataset, concentrating exclusively on the most pertinent proteins for our molecular docking experiments.

The filters applied were as follows:

- **Keyword Filter (Filter 1):** Search terms such as “glucose” were used to identify relevant proteins in the databases.
- **Glucose Filter (Filter 2):** The chemical formula “ C_6_H_12_O_6_” was again employed to identify unique proteins.
- **Homology filter (Filter 3):** Proteins with more than 30% amino acid sequence similarity, indicating homology, were identified through a BLAST search and subsequently removed from the dataset [27,28].

### 2.3 Molecular dynamics simulation studies of docked complexes

The simulation setup of docked complexes was prepared by the CHARMM-GUI solution builder [29]. Details of the simulation setup are given in Table S1. Equilibration and production were conducted under constant volume, pressure, and temperature (1 atm and 310 K) conditions using NAMD 2.14 with CHARMM36m force field[30] for the 100ns. Structural changes during the simulation were measured using root mean square deviation (RMSD), with large RMSD values indicating significant motion and low RMSD values indicating a stable system. Analysis and visualization were performed using VMD [31].

### 2.4 SMD-BD-FDT analysis of docked complexes

Steered molecular dynamics (SMD) simulations were conducted using the Brownian dynamics fluctuation-dissipation theorem (BD-FDT) method [32] to accurately determine the free-energy profiles for glucose unbinding from protein-ligand complexes. The most stable structures of protein-ligand complexes were selected from MD simulations. CHARMM36m was utilized as the preferred force field, with a damping coefficient maintained at 5/ps. The vector connecting the center of mass of the glucose molecule to the center of mass of the alpha carbon atoms (within the binding site residues) was chosen as the pulling direction. The pulling speed was set at 0.002 Å/ps, while the rotational and translational motion of the protein-ligand complexes were fixed during the simulation. The distance between the bound and unbound states was divided into equal segments of 1 Angstrom. Ten forward and reverse paths were sampled for each segment as follows: the system was equilibrated for 0.5 ns with glucose held at the segment’s starting point, then glucose was steered 1 Å away from the binding site to sample a forward path. Subsequently, the system was equilibrated for 0.5 ns with glucose held at the segment’s end point. After equilibration, glucose was steered 1 Å towards the binding site to sample a reverse path. This process was repeated ten times for each segment from the bound state to the dissociated state. The potential of mean force (PMF) was then calculated using these paths from equation (1),

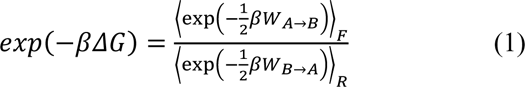

Where the BD-FDT relates the equilibrium free energy to the no equilibrium work according to the equation, here, 𝑊_𝐴→𝐵_ is the force-displacement integral along a forward path from State A to State B, 𝑊_𝐵→𝐴_ is a similarly defined force-displacement integral along a reverse path from State B to State A.

## 3. Results and discussion

### 3.1 Filtering of glucose-binding proteins (GBPs)

A total of 37,325 hits were identified, indicating a potential association with glucose-binding proteins, using Filter 1. Filter 2, which employed the molecular formula of glucose (C_6_H_12_O_6_), narrowed down the protein candidates to 7,919 entries. Subsequently, Filter 3 utilized the CD-HIT algorithm to cluster proteins exhibiting high sequence similarity. This process resulted in a refined set of 1,447 proteins, as detailed in **Table S2.** Follow the link (https://aciitb23.github.io/GBPS.com/) for more information about GBPs.

### 3.2 Virtual screening of glucose-binding proteins (GBPs)

Virtual screening of the selected 1,450 GBPs showed significant variations in their binding affinity with glucose **(Table S3).** Follow the link (https://aciitb23.github.io/GBPS.com/) for more information about GBPs. The GBPs were arranged in decreasing order of their docking scores, Δ𝐸_𝐴𝐷_, reflecting their potential binding affinity. Table 1 lists the docking scores for the top five GPB candidate compounds, along with their respective interacting residues and types of interacting bonds. Whereas Figure 2 shows the three-dimensional structures of these GBPs, Figure 3 shows respective glucose-protein interactions at predicted binding sites. The binding site interactions are shown in greater detail in Figure S1. Among the proteins, 5TA0 exhibited the highest docking score (−7.70 kcal/mol), suggesting a strong binding affinity. For comparison, the commonly used GBP, glucose oxidase (1GAL) was also included in the list. In addition, a GBP with a very low binding affinity (−4.9 kcal/mol), human glycolipid transfer protein (2EVL), was included as a negative control.

**Table 1:**
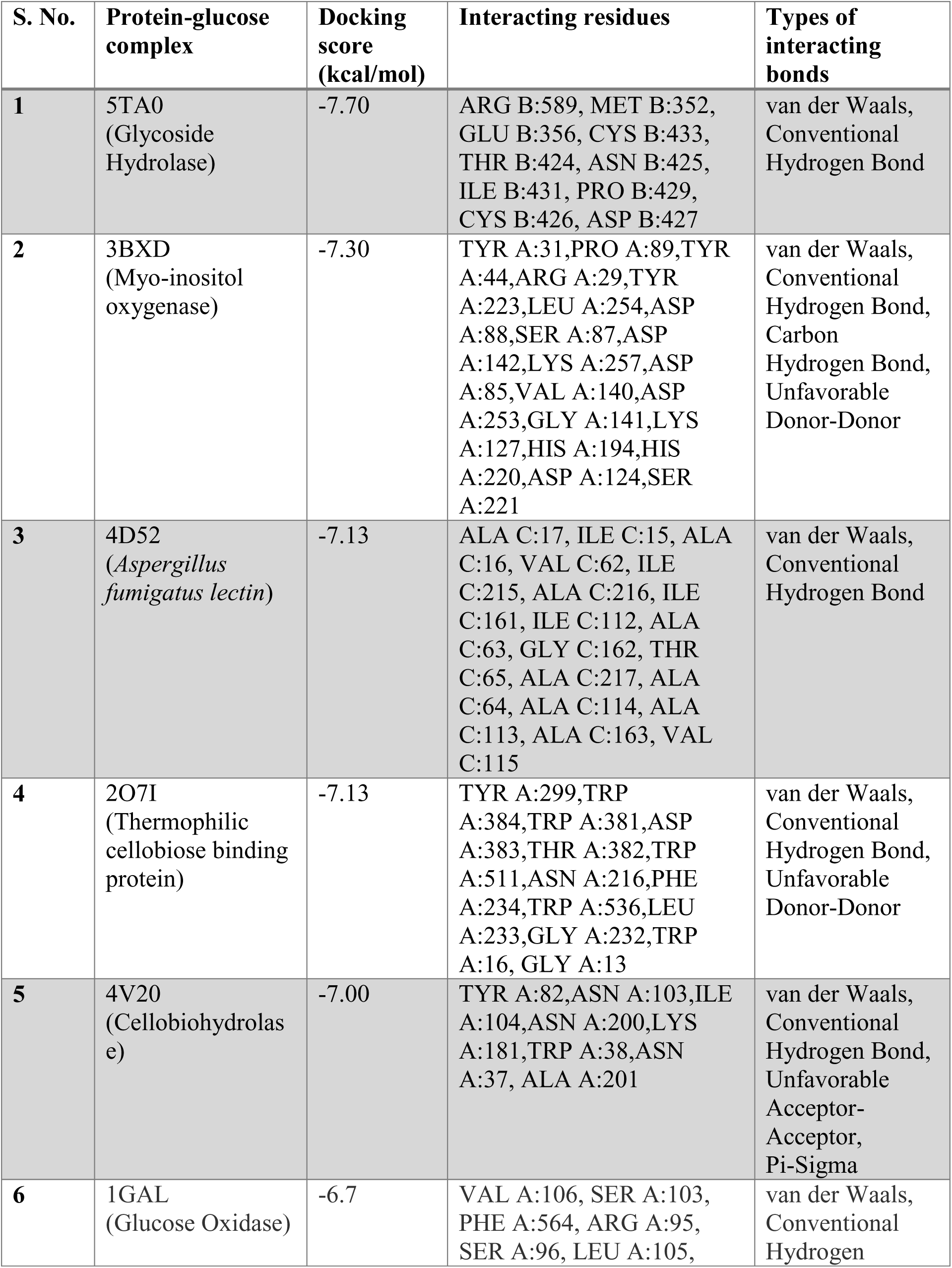
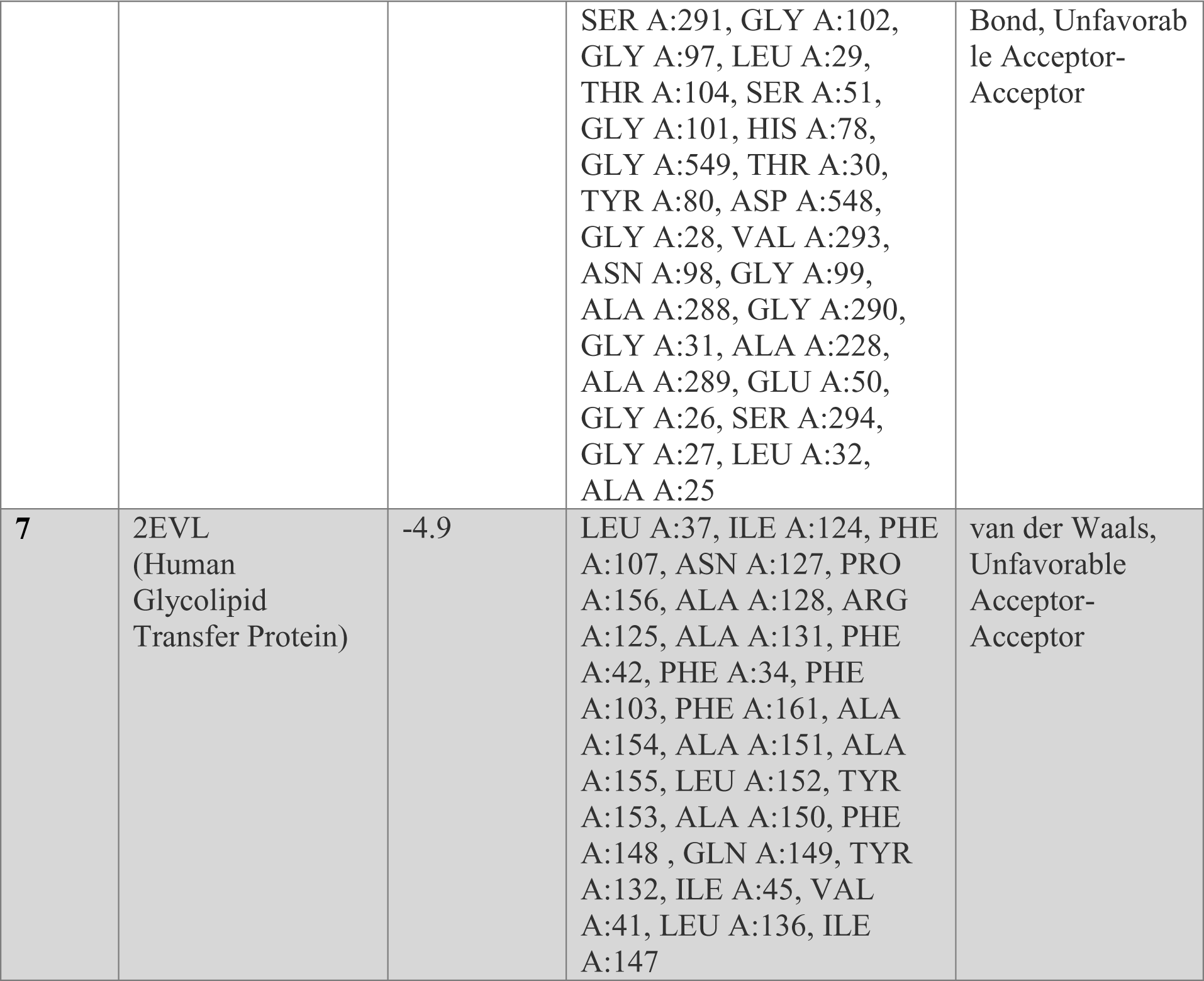
A comprehensive overview of seven ligand-protein complexes, with essential information such as, their respective PDB IDs with names, interacting residues involved in the glucose binding process, and the types of bonds formed between the ligands and proteins.

| S. No. | Protein-glucose complex | Docking score (kcal/mol) | Interacting residues | Types of interacting bonds |
| --- | --- | --- | --- | --- |
| 1 | 5TA0<br>(Glycoside Hydrolase) | -7.70 | ARG B:589, MET B:352, GLU B:356, CYS B:433, THR B:424, ASN B:425, ILE B:431, PRO B:429, CYS B:426, ASP B:427 | van der Waals, Conventional Hydrogen Bond |
| 2 | 3BXD<br>(Myo-inositol oxygenase) | -7.30 | TYR A:31,PRO A:89,TYR A:44,ARG A:29,TYR A:223,LEU A:254,ASP A:88,SER A:87,ASP A:142,LYS A:257,ASP A:85,VAL A:140,ASP A:253,GLY A:141,LYS A:127,HIS A:194,HIS A:220,ASP A:124,SER A:221 | van der Waals, Conventional Hydrogen Bond, Carbon Hydrogen Bond, Unfavorable Donor-Donor |
| 3 | 4D52<br>(*Aspergillus fumigatus* lectin) | -7.13 | ALA C:17, ILE C:15, ALA C:16, VAL C:62, ILE C:215, ALA C:216, ILE C:161, ILE C:112, ALA C:63, GLY C:162, THR C:65, ALA C:217, ALA C:64, ALA C:114, ALA C:113, ALA C:163, VAL C:115 | van der Waals, Conventional Hydrogen Bond |
| 4 | 2O7I<br>(Thermophilic cellobiose binding protein) | -7.13 | TYR A:299,TRP A:384,TRP A:381,ASP A:383,THR A:382,TRP A:511,ASN A:216,PHE A:234,TRP A:536,LEU A:233,GLY A:232,TRP A:16, GLY A:13 | van der Waals, Conventional Hydrogen Bond, Unfavorable Donor-Donor |
| 5 | 4V20<br>(Cellobiohydrolase) | -7.00 | TYR A:82,ASN A:103,ILE A:104,ASN A:200,LYS A:181,TRP A:38,ASN A:37, ALA A:201 | van der Waals, Conventional Hydrogen Bond, Unfavorable Acceptor-Acceptor, Pi-Sigma |
| 6 | 1GAL<br>(Glucose Oxidase) | -6.7 | VAL A:106, SER A:103, PHE A:564, ARG A:95, SER A:96, LEU A:105, | van der Waals, Conventional Hydrogen |
|  |  |  | SER A:291, GLY A:102, GLY A:97, LEU A:29, THR A:104, SER A:51, GLY A:101, HIS A:78, GLY A:549, THR A:30, TYR A:80, ASP A:548, GLY A:28, VAL A:293, ASN A:98, GLY A:99, ALA A:288, GLY A:290, GLY A:31, ALA A:228, ALA A:289, GLU A:50, GLY A:26, SER A:294, GLY A:27, LEU A:32, ALA A:25 | Bond, Unfavorable Acceptor-Acceptor |
| 7 | 2EVL<br>(Human Glycolipid Transfer Protein) | -4.9 | LEU A:37, ILE A:124, PHE A:107, ASN A:127, PRO A:156, ALA A:128, ARG A:125, ALA A:131, PHE A:42, PHE A:34, PHE A:103, PHE A:161, ALA A:154, ALA A:151, ALA A:155, LEU A:152, TYR A:153, ALA A:150, PHE A:148, GLN A:149, TYR A:132, ILE A:45, VAL A:41, LEU A:136, ILE A:147 | van der Waals, Unfavorable Acceptor-Acceptor |

**Figure 2:**
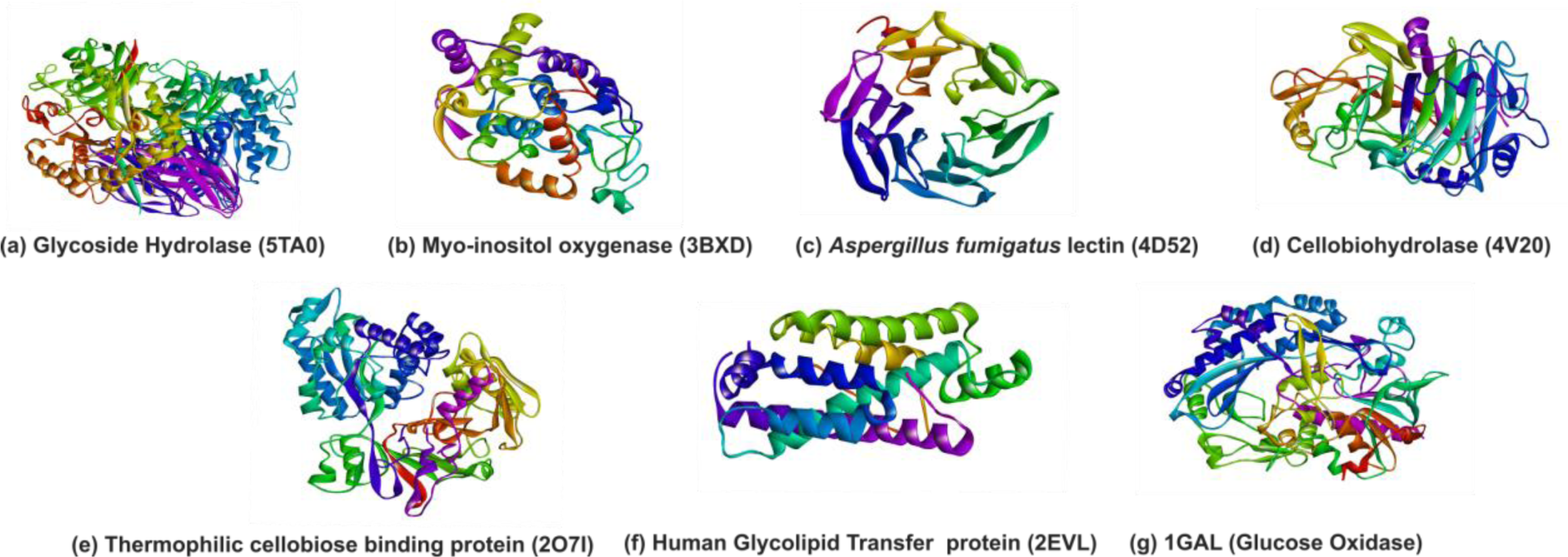
Glucose-binding proteins (GBPs) identified through molecular docking screening. The proteins are labelled using their respective PDB IDs: (a) Glycoside Hydrolase (5TA0), (b) Myo-inositol oxygenase (3BXD), (c) *Aspergillus fumigatus* lectin (4D52), (d) Cellobiohydrolase (4V20), (e) Thermophilic cellobiose binding protein (2O7I), (f) Human Glycolipid Transfer Protein (2EVL), and (g) Glucose Oxidase (1GAL).

**Figure 3:**
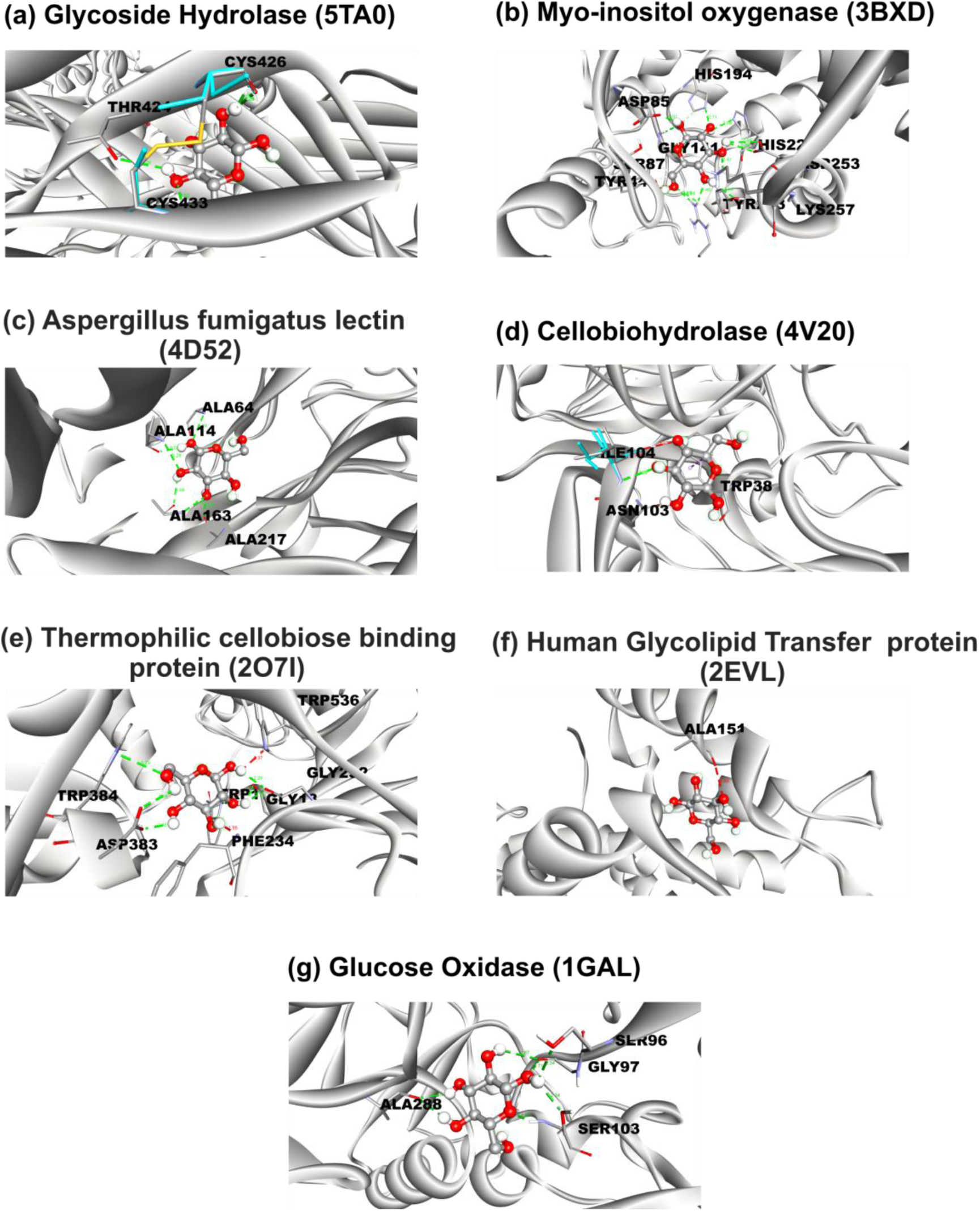
Visualization of intermolecular interactions of the glucose molecule bound to, (a) Glycoside Hydrolase (5TA0), (b) Myo-inositol oxygenase (3BXD), (c) *Aspergillus fumigatus* lectin (4D52), (d) Cellobiohydrolase (4V20), (e) Thermophilic cellobiose binding protein (2O7I), (f) Human Glycolipid Transfer Protein (2EVL), and (g) Glucose Oxidase (1GAL). Dashed lines indicate various interaction types, with each color representing a specific interaction type (*green* – hydrogen bond, *red* – Unfavorable Acceptor-Acceptor). Residues are depicted with their three-letter codes and residue numbers.

### 3.3 Simulation of docked complexes

The stability of the docked complexes (candidate protein + glucose) and associated structural changes were monitored with 100 ns long MD simulations (Figure 4a). RMSD values for all docked complexes were found to vary between 1 – 6 (Å). RMSD values for the 4D52 complex were the most stable among the seven GBPs (Figures 4b, c). In addition, 4V20 and 3BXD complexes also demonstrated higher stability compared to the rest of the complexes. The 2O7I complex exhibited relatively higher RMSD values compared to the other docked complexes. Hydrogen bond occupancy was calculated for all protein-ligand complexes, revealing varying degrees of hydrogen bond occupancy for each complex (Figure S2). These results suggested that RMSD values alone may not be the primary determinant of GBP complex stability.

**Figure 4:**
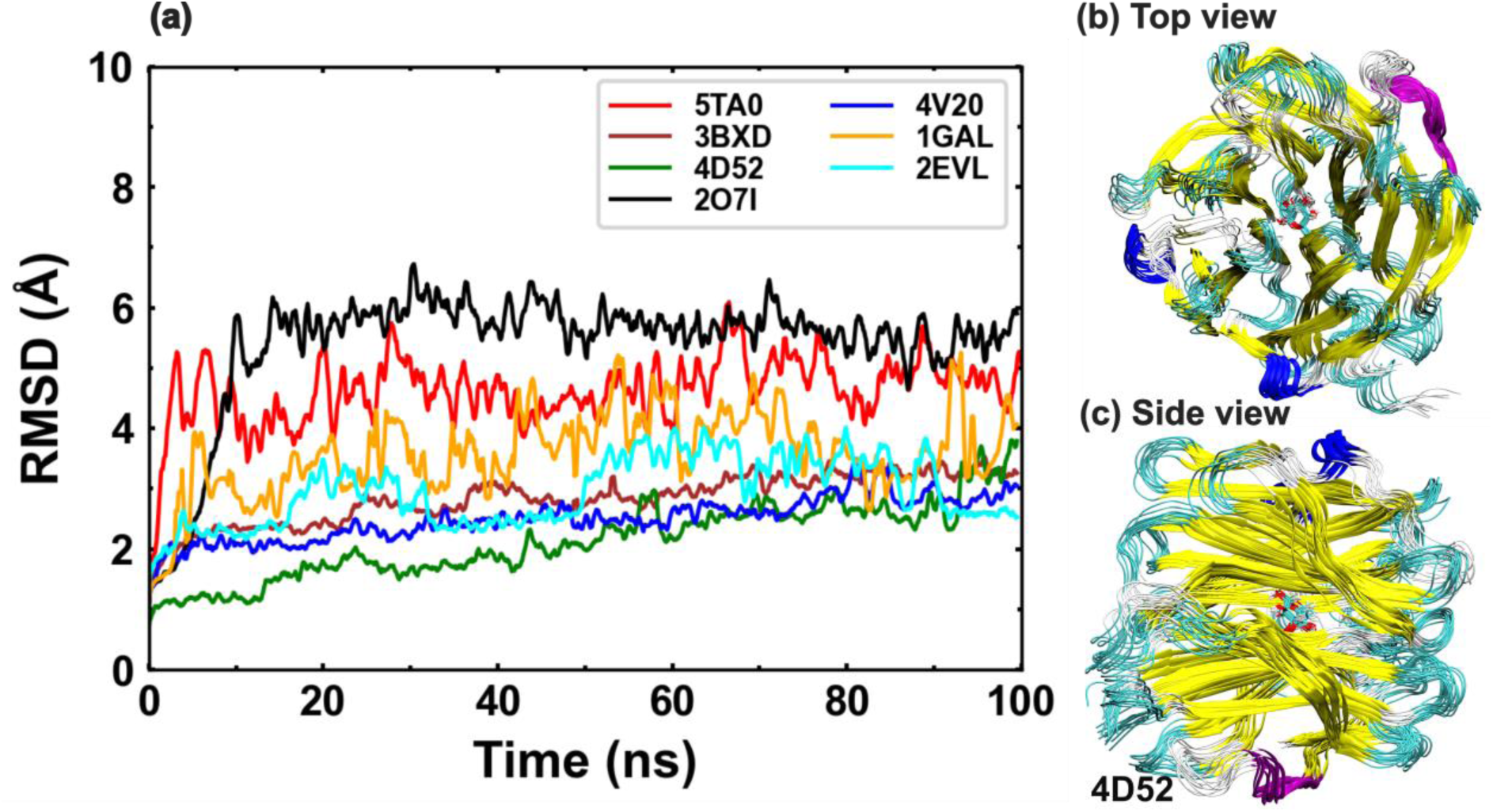
**(a)** Comparative RMSD analysis of the docked protein-glucose complexes from MD simulations. **(b, c)** Different conformations of the 4D52 protein-glucose complex observed over the simulation, showing the glucose molecule (licorice/stick representation) and the 4D52 protein (ribbon/thread-like representation).

However, not all complexes remained stable during the simulations. The distance of the glucose molecule from the binding sites (CA atom) of GBPs, 𝑑_𝐺_, was used as a measure of the integrity of the GBP-glucose complex. Plots of variation of 𝑑_𝐺_ with time are shown in Figures 5a and b. A residence time, 𝜏_𝑏_, for a specific GBP was defined as the time after which the corresponding 𝑑_𝐺_ value started diverging significantly from its initial value. Based on 𝜏_𝑏_ values (estimated from the 𝑑_𝐺_ versus time plots), the seven GBPs were characterized as candidates with either strong (Figure 5a) or weak (Figure 5b) binding affinities. Whereas 4D52 and 1GAL demonstrated strong binding affinities, 4V20, 2O7I, and 2EVL showed the lowest binding with glucose. Complexes of 3BXD and 5TA0 remained bound to glucose for approximately 35 ns and 70 ns, respectively. Since these complexes remained stable for much longer when compared to the weakly-binding GBPs, they were also classified as strongly-binding GBPs. The residence time, estimated from these calculations, was used as a parameter in the selection process because it describes the dynamics of ligand binding to the GBP. Glucose binding/unbinding times of a strongly-binding (5TA0) and a weakly-binding (2EVL) GBPs are shown in Figures 5c and d, respectively.

**Figure 5:**
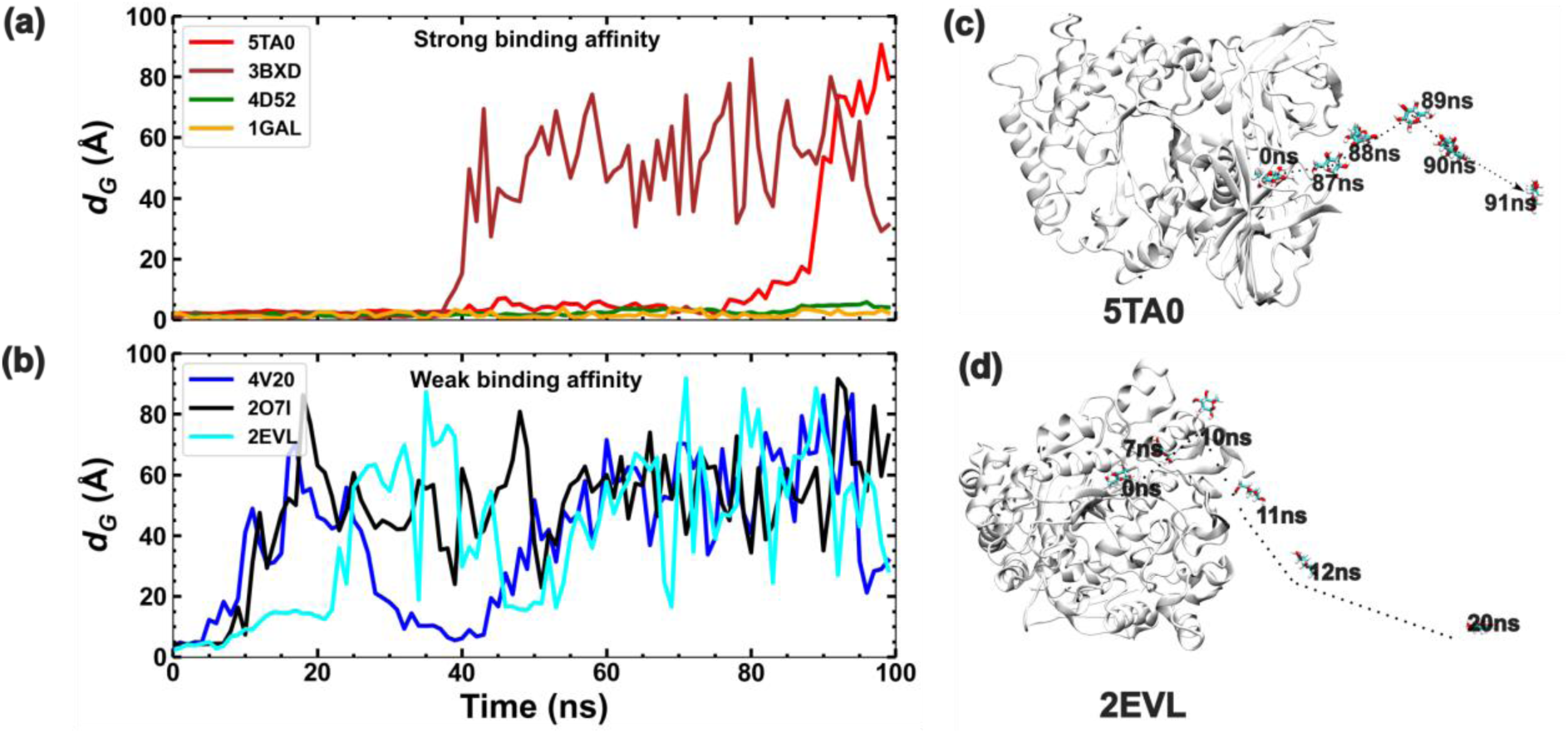
**(a-b)** Plots showing the variation of distance of the glucose molecule from the binding sites (CA atom) of GBPs, 𝑑_𝐺_, as a function of time for all seven GBPs. Unbinding paths of the glucose molecule for the cases of (c) 5TA0 and (d) 2EVL, showing the large difference in their relative stabilities with respect to glucose binding.

### 3.4 Potential of mean force from SMD-BD-FDT

Using the SMD-BD-FDT simulations, the potential of mean force (PMF) curves for the seven GBPs, were estimated as a function of the distance of the ligand from the binding site, 𝜉 (Figure 6). The binding free energy of a ligand-protein complex was inferred from the PMF curve as the depth of the potential well. Most of the BD-FDT binding energy values, Δ𝐺_𝑏_, were different from the binding energies estimated from the AutoDock values (Figure 7). For instance, BD-FDT predicted a binding energy of −16.7 kcal/mol for 5TAO, which was significantly higher than the value estimated by AutoDock at −7.7 kcal/mol. Interestingly, 2EVL, which was selected for its low binding score, displayed a low binding energy value of −2.6 kcal/mol in the BD-FDT simulations as well. Moreover, 2O7I demonstrated a binding free energy of −11.5 kcal/mol, higher than the docking prediction of −7.13 kcal/mol. However, the PMF curve showed a negative curvature, indicating weak binding of glucose with the protein. This inference is consistent with the extremely short 𝜏_𝑏_ for 2O7I (Figure 5c) and in conflict with the prediction of a stable glucose-2O7I complex from AutoDock. It shows that AutoDock scores for binding energies may not capture all dynamical aspects of ligand binding to the GBP. Hence, AutoDock predictions need to be guided by more rigorous physics-based simulations to generate rules for protein selection. Representative paths corresponding to glucose unbinding for each GBP are shown in Figure S3.

**Figure 6:**
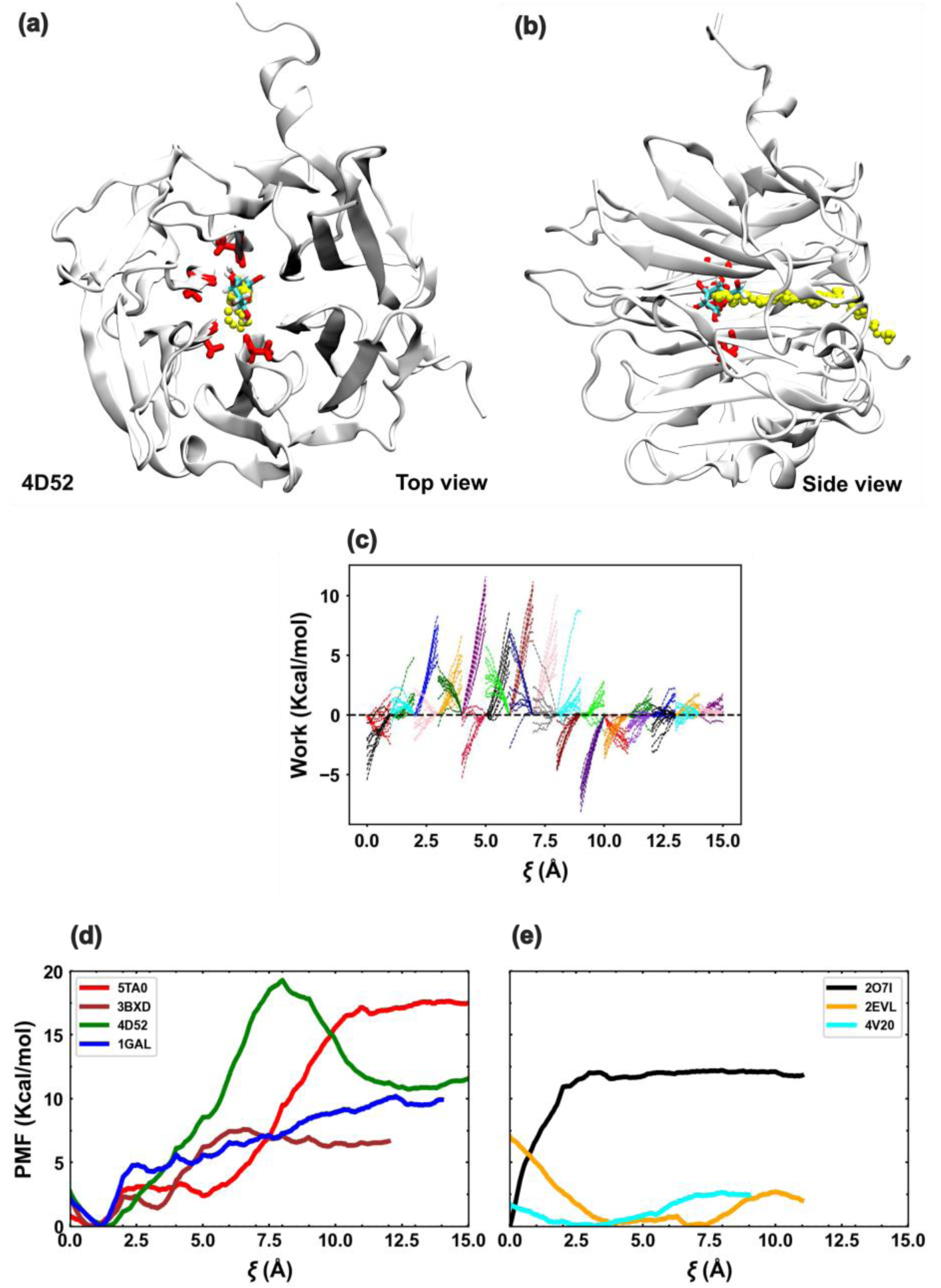
(a) Top and (b) side views of the 4D52 protein with the glucose binding site and bound glucose. The glucose binding site is shown with thick red lines protruding from the protein, and the glucose is shown in *red, white,* and *cyan* lines. The unbinding path is shown by yellow beads in both (a) and (b). **(c)** Work done along the forward (spreading out towards right) and reverse (spreading out towards left) pulling paths in the BD-FDT SMD simulations **(c-d)** PMF profiles of glucose unbinding from the GBPs estimated from SMD simulations.

**Figure 7:**
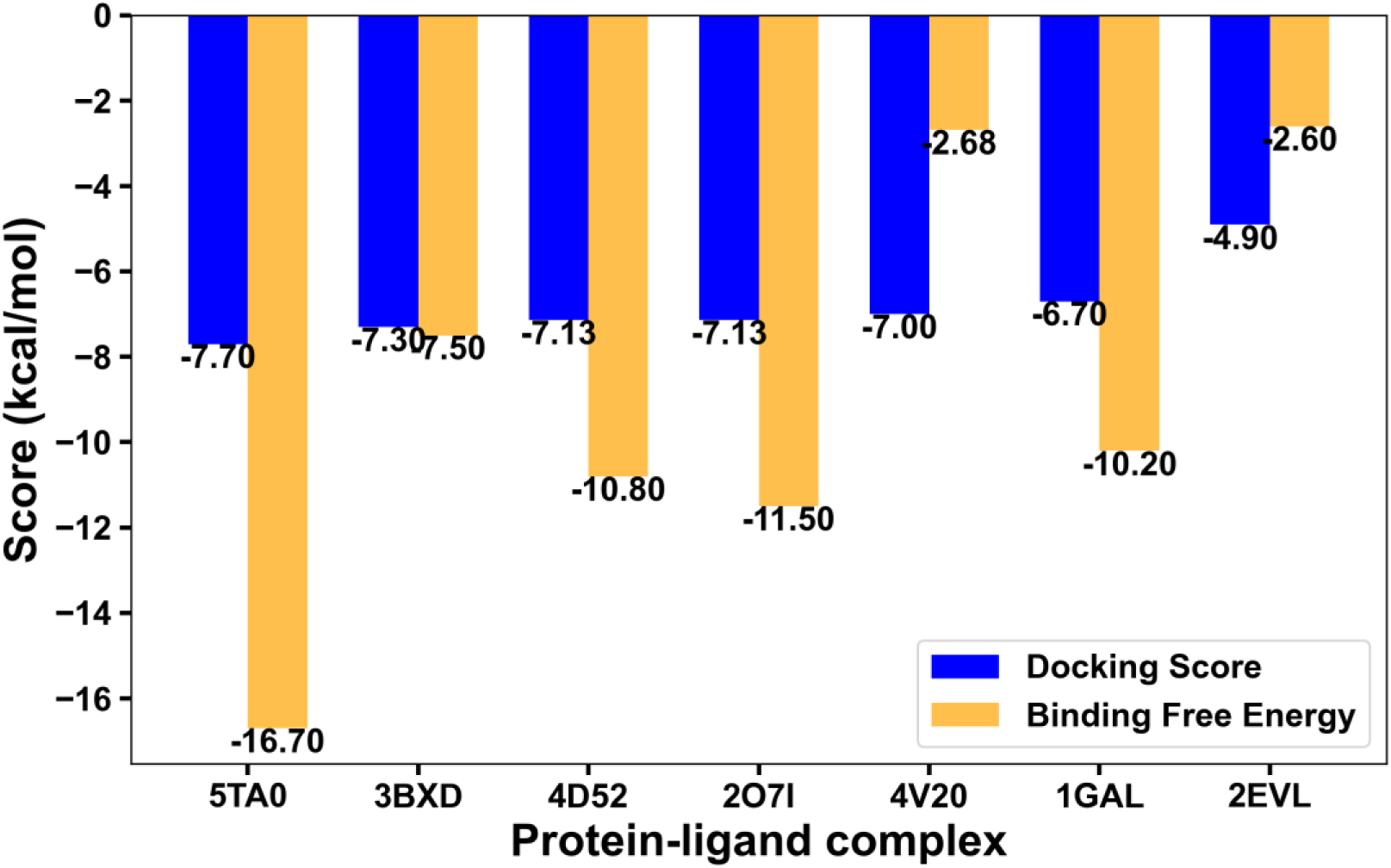
A comparison of GBP docking scores (from molecular docking) and binding free energies of protein-ligand complexes (obtained from BD-FDT SMD simulations).

The examination of individual cases further emphasized the diverse characteristics of the complexes. The analysis of the potential of 4V20 elucidated its limited binding capacity, supported by a shallow well and subsequent dissociation, confirmed by a low PMF value. Similarly, 2EVL showed weakened bonding, as evidenced by the shallow well and subsequent unbinding of glucose beyond a certain threshold. On the other hand, the unbinding pathways of 2O7I and 1GAL revealed distinct patterns, reflecting the complexities of their interactions. While 2O7I demonstrated a relatively abrupt dissociation, 1GAL displayed an intermediate state during the process, ultimately leading to complete detachment. Notably, 5TA0 exhibited unique unbinding patterns characterized by distinct potential wells and intermediate states, underscoring the intricate nature of their binding mechanisms. Figure 7 compares binding energy predictions from docking, Δ𝐸_𝐴𝐷_, with those from SMD simulations, Δ𝐺_𝑏_. Both Δ𝐸_𝐴𝐷_and Δ𝐺_𝑏_values were the highest for 5TA0. Other proteins with high Δ𝐸_𝐴𝐷_ scores (3BXD, 4D52, and 1GAL) also demonstrated high Δ𝐺_𝑏_ values. For 2EVL, a weakly-binding GBP, both Δ𝐸_𝐴𝐷_and Δ𝐺_𝑏_values were low. These observations indicated a relatively strong correlation between Δ𝐸_𝐴𝐷_and Δ𝐺_𝑏_values. They also set the stage for finding other correlations between energy-minimization-based AutoDock predictions and more detailed atomistic MD simulations. Such correlations can assist in developing a rational GBP selection framework based on relatively quicker, albeit approximate, docking simulations.

### 3.5 Correlating docking predictions with MD simulations

A comprehensive analysis was carried out to correlate results from docking with those from MD simulations. It should be noted that absolute values of the binding energies, |Δ𝐺_𝑏_| and |Δ𝐸_𝐴𝐷,_| are represented in these plots. Key parameters derived from MD simulations, including glucose residence time (𝜏_𝑏_), glucose binding free energy (|Δ𝐺_𝑏_|), and the hydrogen bond occupancy (𝑛_𝐻𝐵𝐷_), were selected as the first set of parameters. The second set of parameters, derived from docking simulations, included the docking energy (|Δ𝐸_𝐴𝐷_|), cavity depth of the binding site from the GBP surface (𝑑_𝑐_), and the number of hydrogen bonds between glucose and GBP residues (𝑛_𝐴𝐷_). Correlations were sought between parameters in the first set with those in the second set. Figures 8a – c plot 𝜏_𝑏_ as a function of the docking parameters from the second set. Similarly, Figures 8d – f plot |Δ𝐺_𝑏_| as a function of the second set docking parameters.

**Figure 8:**
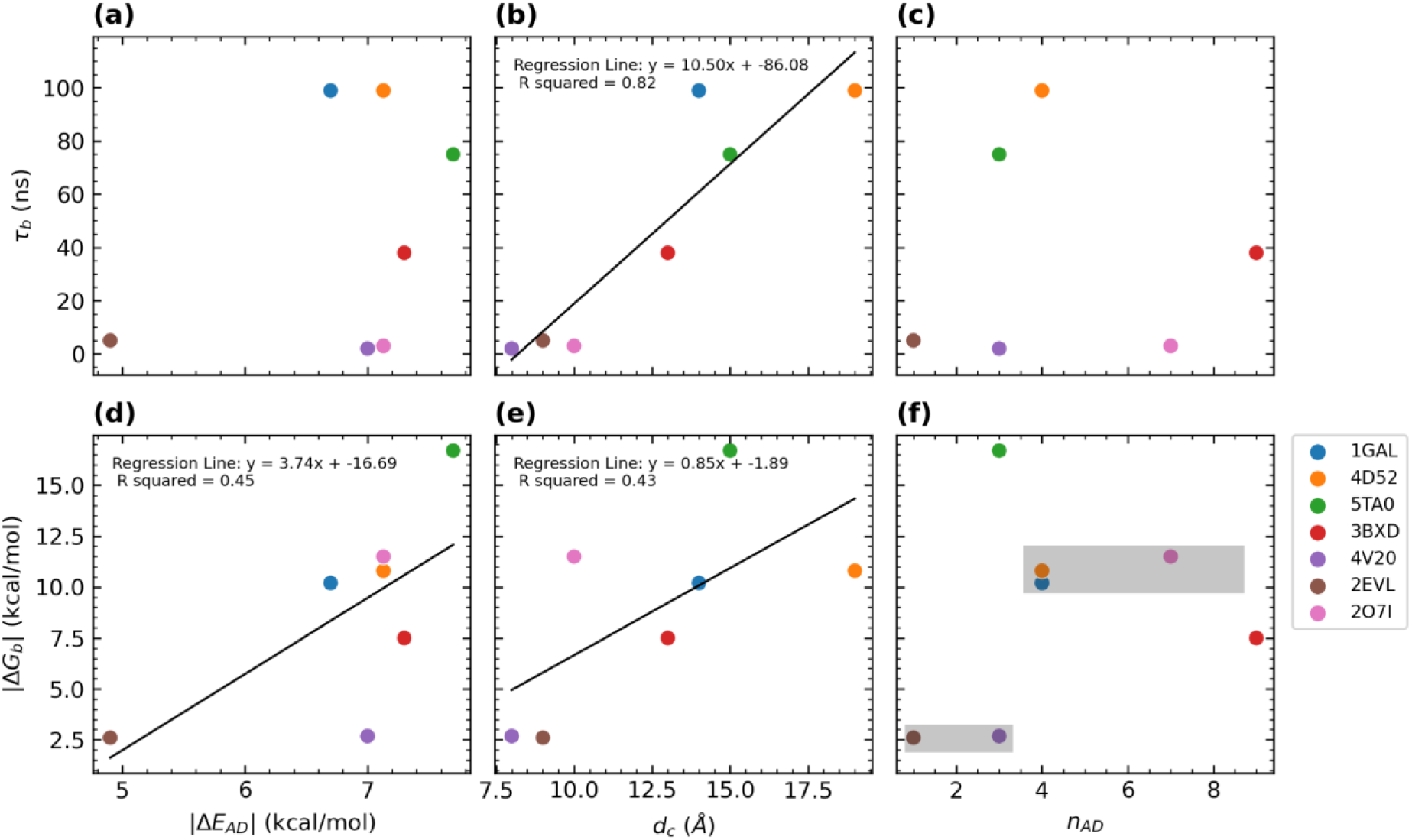
Correlation plots between parameters from two different sets, (i) ligand-binding (𝜏_𝑏_ and |Δ𝐺_𝑏_| from MD simulations) and (b) binding parameters (|Δ𝐸_𝐴𝐷_|, 𝑑_𝑐_, and 𝑛_𝐴𝐷_ from docking simulations).

Whereas 𝜏_𝑏_did not show any correlation with either |Δ𝐸_𝐴𝐷_| or 𝑛_𝐴𝐷_ (Figures 8a,c), it showed a strong linear correlation with 𝑑_𝑐_ (Figure 8b). The correlation plot of 𝜏_𝑏_vs 𝑑_𝑐_showed that a large cavity depth resulted in a strongly-bound and stable glucose-GBP complexes. A comparison with MD results from Figures 5a and b indicated that weakly-bound GBPs (2EVL, 4V20, and 2O7I) had 𝑑_𝑐_ < 10Å. Interestingly, both 4V20 and 2O7I had high |Δ𝐸_𝐴𝐷_| values, implying that a high docking score may not necessarily lead to stable glucose-GBP complexes.

The binding free energy, |Δ𝐺_𝑏_|, showed stronger correlations with the three docking parameters (Figures 8d – f). It increased almost linearly with increasing value of both |Δ𝐸_𝐴𝐷_| (Figure 8d), but there were a couple of anomalies. As stated above, 4V20 had a high |Δ𝐸_𝐴𝐷_| but did not form a stable complex (Figure 5b) because of which it had a low |Δ𝐺_𝑏_| value. Similarly, 2O7I also formed an unstable complex (though it had a high |Δ𝐸_𝐴𝐷_|, but a high |Δ𝐺_𝑏_| value was interpreted from its anomalous PMF curve in Figure 6e. Again, |Δ𝐺_𝑏_| showed a linear correlation with 𝑑_𝑐_ (Figure 8e). In general, GBPs with deeper cavities resulted in higher |Δ𝐺_𝑏_| values. Thus, cavity depth emerged as a reliable indicator of glucose binding with a GBP. The correlation plots in Figures 8b,e suggested that stable glucose-GBP complexes emerged for 𝑑_𝑐_ > 12Å. Plotting the variation of |Δ𝐺_𝑏_| with respect to 𝑛_𝐴𝐷_ provided insight into the effect of the bonding environment at the ligand binding site on the stability of the glucose-GBP complex. A sigmoidal dependence of |Δ𝐺_𝑏_| with respect to 𝑛_𝐴𝐷_was inferred from Figure 8f. Whereas GBPs with 𝑛𝐴𝐷≤3 corresponded to low Δ𝐺𝑏 values (weakly binding), GBPs with 𝑛_𝐴𝐷_ > 3 corresponded to stable complexes with Δ𝐺_𝑏_ values close to 10 kcal/mol. Thus, |Δ𝐺_𝑏_| versus 𝑛_𝐴𝐷_plot indicated a threshold value of 𝑛_𝐴𝐷_ = 3, above which stable complexes could be found.

To validate these correlations interpreted from Figure 8, |Δ𝐺_𝑏_| was plotted with respect to two other parameters also obtained from MD data, namely, 𝑛_𝐻𝐵𝐷_ and 𝜏_𝑏_ (Figure 9). The plot of |Δ𝐺_𝑏_| with respect to 𝑛_𝐻𝐵𝐷_showed a sigmoidal behavior similar to the plot in Figure 8f, indicating that a certain minimum number of hydrogen bonds (at the binding site) are required for stable glucose-GBP complexes. Similarly, low 𝜏_𝑏_ values corresponded to low |Δ𝐺_𝑏_| values and, by extension, a weakly-bound glucose molecule (Figure 9b).

**Figure 9:**
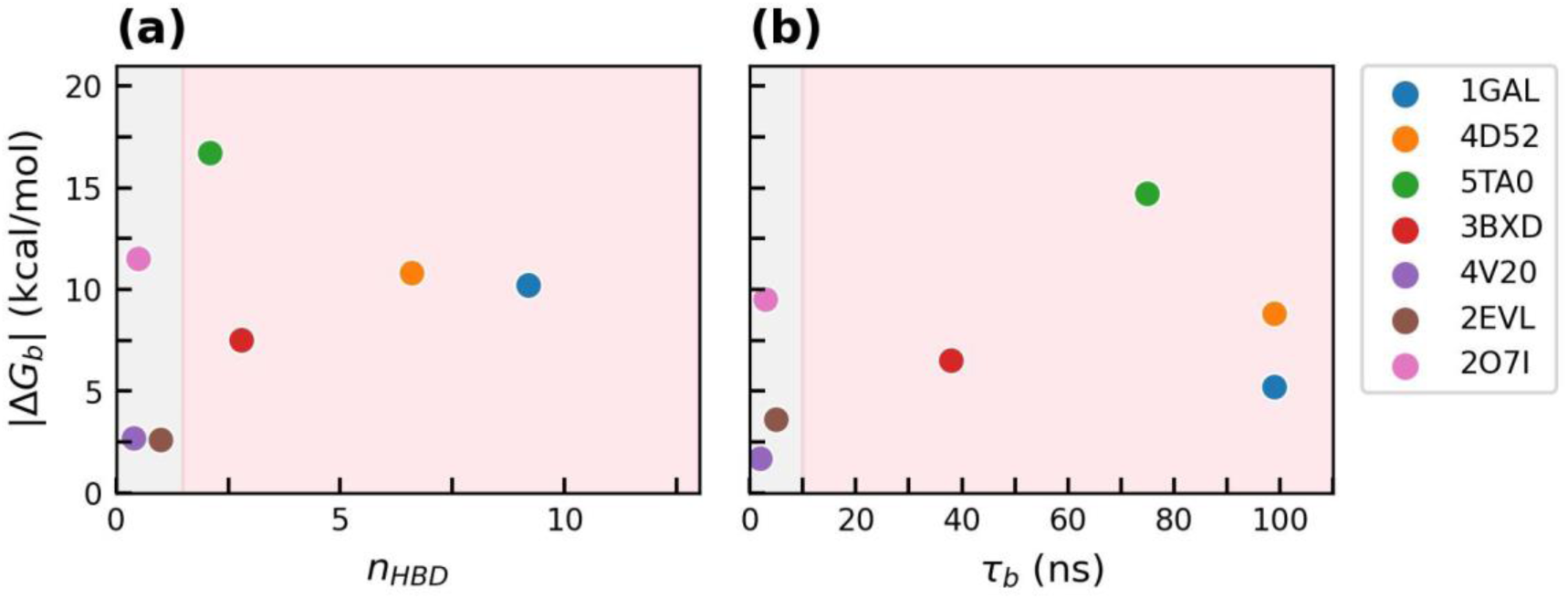
Correlation plot between (a) |Δ𝐺_𝑏_| and 𝑛_𝐻𝐵𝐷_, and (b) |Δ𝐺_𝑏_| and 𝜏_b_ (ns). Whereas, 2O7I, 2EVL, and 2O7I, with lowest 𝑛_𝐻𝐵𝐷_and 𝜏_b_ (ns) values, are shown in light grey color, 1GAL, 4D52, 5TA0, and 3BXD with high HBD occupancy and 𝜏_b_ (ns) values shown in light pink color.

Thus, the correlation plots in Figure 8 helped in arriving at a set of docking parameters that would reliably predict GBPs that bound strongly to glucose. Using these parameters would make the protein selection process more efficient since these parameters would be obtained from relatively faster docking simulations. Based on the above discussion, a combination of threshold values of Δ𝐸𝐴𝐷, 𝑑𝑐, and 𝑛_𝐴𝐷_can be used to predict candidate GBPs reliably. Specifically, a GBP with docking parameters corresponding to |Δ𝐸_𝐴𝐷_ | > 7.0, 𝑑_𝑐_ > 12Å, and 𝑛_𝐴𝐷_ > 3 should result in the formation of a stable glucose-protein complex.

Figure 10 is a flowchart that outlines a “GBP selection pipeline” that incorporates the above-mentioned selection rules (based only on docking data) and a few test MD simulations. It comprises the following steps,

i. Use Filters 1, 2, and 3 to select candidates from the database
ii. Apply threshold values of |Δ𝐸_𝐴𝐷_| > 7.0 kcal/mol, 𝑑_𝑐_ > 12Å and 𝑛_𝐴𝐷_ > 3 to arrive at potentially strongly-binding GBPs
iii. Computational validation of selection process: Conduct short (100 ns) MD simulations on a small sample of proteins (≈10 proteins) to test the stability of the glucose-protein complex (along the lines of Figure 5)
iv. Experimental validation: Use top candidates for further experimental binding studies

**Figure 10:**
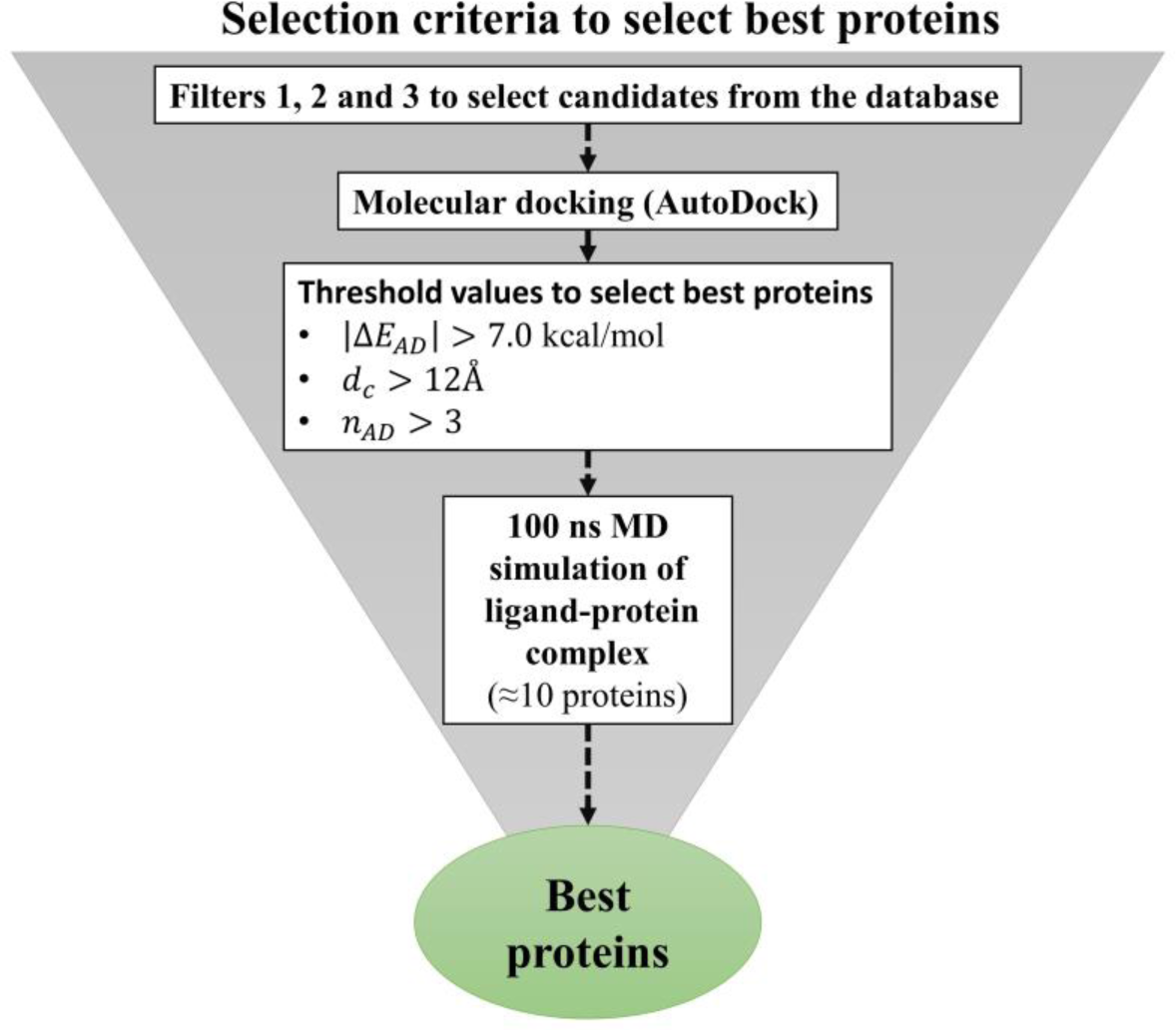
A flowchart showing the proposed GBP selection criteria based on quick docking simulations and only a limited number of relatively faster MD simulations.

## 4. Conclusion

Biosensors are valuable tools for monitoring the quality of fruit juices due to their fast response time, portability, and cost-effectiveness. Enzymatic biosensors have proven successful due to their high specificity and low manufacturing cost. In the current work, unique glucose-binding proteins were identified using a high-throughput bioinformatics pipeline, which included sequence-based analysis, molecular docking, and MD simulations. A total of 37,325 GBP hits were identified from the PDB database, and subsequently, a total of 1,447 unique GBPs were found suitable. Molecular docking data provided approximate values for binding (docking) energies, binding cavity depths, and state of hydrogen bonding at the binding site (Figures 3 and 8). Detailed binding dynamics of a smaller subset of seven GBPs were exhaustively investigated using MD simulations. MD simulations provided estimates of glucose binding times to corresponding GBPs (Figure 5). BD-FDT SMD simulations were used to estimate the binding free energies of the ligand-protein complex (Figure 6).

Several parameters belonging to two different sets, (a) ligand-binding (from MD simulations) and (b) binding parameters (from docking simulations), were compared to extract correlations between them (Figures 7 and 8). The objective was to propose parameter values based only on faster docking simulations that could be used reliably to select GBPs. The correlation plots suggested that the following threshold values of three docking parameters: Δ𝐸𝐴𝐷>7.0 kcal/mol, 𝑑𝑐>12Å, and 𝑛_𝐴𝐷_ > 3 can be used to predict candidate GBPs reliably. All selection criteria were based only on data generated from quick docking simulations. A high-throughput and accurate protein selection process based on quick docking simulations was proposed to screen GBPs for glucose biosensing (Figure 10).

Top candidates from the screening process can be tested in laboratory experiments to study glucose binding. This would constitute a test for the validity of the proposed scheme. Experimental validation was not in the scope of the current work. However, 1GAL (a widely used GBP) met all three proposed selection criteria. The current scheme can also be extended to other ligand-binding proteins for different analytes present in food processing, including fruit juices.

## Acknowledgments

The authors acknowledge support in the form of access to the Spacetime High-Performance Computing (HPC) resource at IIT Bombay.

## Author’s Contributions

A.C., P.D., and A.S.P. designed the research. A.C. developed a high throughput screening pipeline for screening GBP candidates. A.C. and A.M. prepared the initial manuscript draft.

A.M. performed SMD simulations, analyzed data, and created figures. P.D. and D.D. provided inputs on glucose sensing and analysis. A.C. and A.S.P. contributed to the writing and editing of the final manuscript.

## Data availability

Follow the link (https://aciitb23.github.io/GBPS.com/) for more information about screened GBPs.

