## Supplemental Table S1 and Figure S1, S2, S3 for "High-throughput selection of glucose-binding proteins from massive datasets: Integrating molecular docking and molecular dynamics simulations"

**Table S1:** Simulation setup details of Glucose-binding proteins.

| S.No. | Protein-glucose complex | Total no. of atoms | System size (X=Y=Z) | Na+ ions | $Cl^-$ ions | Water | Protein | Time (ns) |
|---|---|---|---|---|---|---|---|---|
| 1. | 5TA0 (Glycoside Hydrolase) | 2,69,544 | 142 | 242 | 236 | 249090 | 19952 | 100 ns |
| 2. | 3BXD (myo-inositol oxygenase) | 44,293 | 78 | 50 | 38 | 40047 | 4134 | 100 ns |
| 3. | 4D52 (*Aspergillus fumigatus lectin*) | 2,87,242 | 145 | 254 | 254 | 267663 | 19047 | 100 ns |
| 4. | 4V20 (Cellobiohydrolase) | 61,598 | 87 | 71 | 52 | 55122 | 6329 | 100 ns |
| 5. | 2O7I (thermophilic cellobiose binding protein) | 85,590 | 97 | 91 | 72 | 76026 | 9377 | 100 ns |
| 6. | 2EVL (Glycolipid transfer protein) | 48,152 | 78 | 42 | 42 | 44703 | 3341 | 100 ns |
| 7. | 1GAL (Glucose oxidase) | 83,441 | 94 | 99 | 70 | 74526 | 8722 | 100 ns |

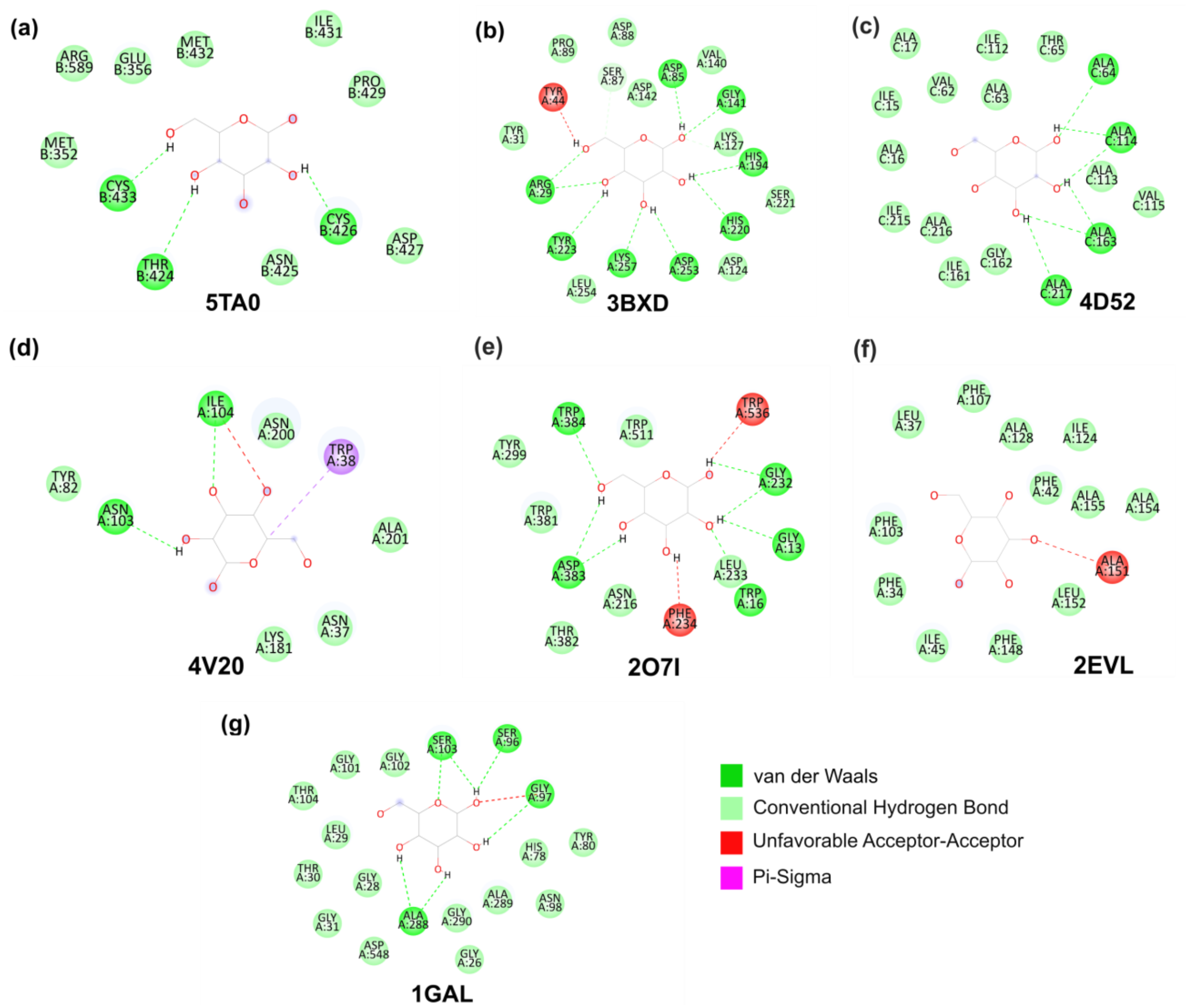


**Figure S1:** 2D representation of intermolecular interactions of the glucose molecule bound to, (a) Glycoside Hydrolase (5TA0), (b) Myo-inositol oxygenase (3BXD), (c) *Aspergillus fumigatus* lectin (4D52), (d) Cellobiohydrolase (4V20), (e) Thermophilic cellobiose binding protein (2O7I), (f) Human Glycolipid Transfer Protein (2EVL), and (g) Glucose Oxidase (1GAL). Dashed lines indicate various interaction types, with each color representing a specific interaction type (*green* – hydrogen bond, *red* – Unfavorable Acceptor-Acceptor). Residues are depicted with their three-letter codes and residue numbers.

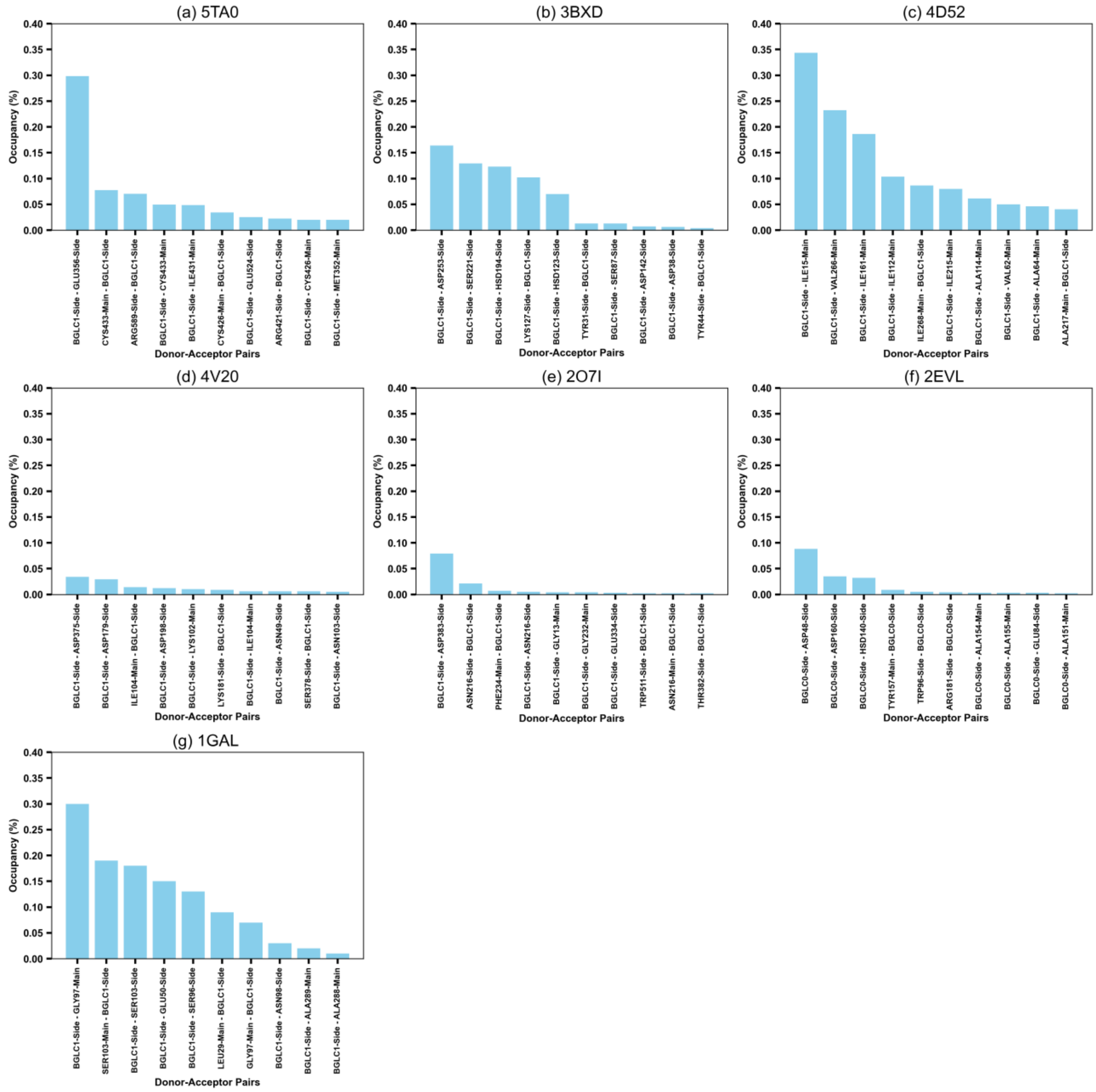


**Figure S2:** Bar plot represents top 10 $n_{HBD}$ in seven ligand-protein complexes. (a) Glycoside Hydrolase (5TA0), (b) Myo-inositol oxygenase (3BXD), (c) *Aspergillus fumigatus* lectin (4D52), (d) Cellobiohydrolase (4V20), (e) Thermophilic cellobiose binding protein (2O7I), (f) Human Glycolipid Transfer Protein (2EVL), and (g) Glucose Oxidase (1GAL).

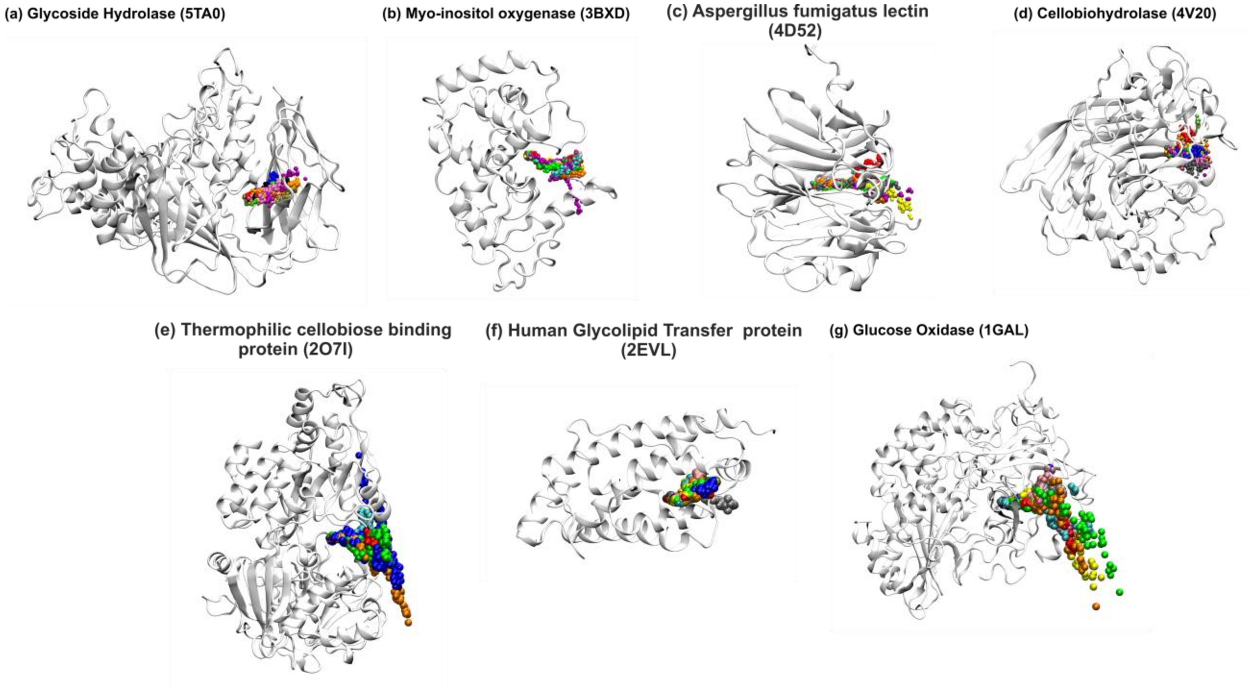


**Figure S3:** Ligand path ($d_G$) of 10 trajectories represented by beads (Unbinding from Left to Right direction for all proteins) in different colors. (a) Glucose Oxidase (1GAL), (b) Human Glycolipid Transfer Protein (2EVL), (c) Thermophilic cellobiose binding protein (2O7I), (d) Myo-inositol oxygenase (3BXD), (e) *Aspergillus fumigatus* lectin (4D52), (f) Cellobiohydrolase (4V20) and (g) Glycoside Hydrolase (5TA0).
